# Transcriptomic responses to within- and transgenerational environmental warming in a cold-adapted salmonid

**DOI:** 10.1101/2022.10.21.513272

**Authors:** Chantelle M. Penney, Gary Burness, Gerardo Zapata, François Lefebvre, Chris C. Wilson

## Abstract

Cold-adapted species are particularly threatened by climate change as rates of environmental warming outpace the ability of many populations adapt. Recent evidence suggests that transgenerational thermal plasticity may play a role in the response of cold-adapted organisms to long-term changes in temperature. Using RNA sequencing, we explored differential gene expression of lake trout (*Salvelinus namaycush*), a cold-adapted species, to examine the molecular processes that respond to elevated temperatures under conditions of within-generation (offspring) and transgenerational (parental) warm acclimation. We hypothesized that genes associated with metabolism, growth and thermal stress/tolerance would be differentially expressed in juvenile lake trout offspring depending on their own acclimation temperature and that of their parents. While parental warm acclimation did have a transgenerational effect on gene expression in their offspring, within-generation (offspring) warm acclimation had a larger effect on the number of differentially expressed genes. Differentially expressed genes enriched pathways for thermal stress, signaling processes, immune function, and transcription regulation and depended on the acclimation temperature of the offspring in isolation or in combination with parental warm acclimation. We provide evidence of the transgenerational response to warming at the transcriptional level in lake trout, which should be useful for future studies of transcriptomics and plasticity in this and other cold-adapted species.

## Introduction

Temperate and Arctic regions are warming rapidly; surface temperatures in Canada have increased by ~1.7°C in under a century with even greater warming happening in the Arctic (~2.3°C; Zhang et al. 2019). As a result, high-latitude and cold-adapted organisms are experiencing, and will continue to experience, rapid temperature increases that may threaten their survival (Beitinger and Bennett 2000; Burkhead 2012; IPCC 2021). Range expansion to track suitable conditions is particularly challenging for species and populations with limited dispersal options, such as many freshwater fishes (Guzzo and Blanchfield 2017; Smith et al. 2020). Warming rates may outpace adaptive responses (Visser 2008; Crozier and Hutchings 2014; Comte and Olden 2017), particularly for those with low standing genetic variation (Willi et al. 2006; Wilson 2017). In the absence of standing genetic variation for coordinated adaptive responses throughout core metabolic pathways, population responses would be limited to plastic thermal acclimation, but the low physiological plasticity typical of many cold-adapted species means they likely will be particularly vulnerable to rapidly changing environments (Somero 2010; Kelly et al. 2014; Kelly et al. 2018). Individuals that can withstand warming may have to rely heavily on energetic resources that would normally support growth or reproduction, thus having potentially negative downstream effects on the population (Somero 2010; Rosenfeld et al. 2020). A lack of adaptive genetic variation, low capacity for temperature acclimation, and rapid environmental change are predicted to result in widespread species extirpations (Bennett et al. 2019; Morash et al. 2021).

Species may be able to buffer the negative impact of environmental warming to some extent through transgenerational acclimation, also known as transgenerational plasticity (TGP). As the name suggests, TGP refers to the phenotypic changes that occur over the course of multiple generations (Jablonka et al. 1992; Bell and Hellmann 2019; Bonduriansky 2021). This involves non-genetic inheritance of environmentally-driven changes, including parental effects, where the experiences of one generation are reflected in the morphology, physiology or behaviour of a subsequent generation (Hellman et al. 2020; Lee et al. 2020; Bonduriansky 2021).

TGP occurs through modulation of gene expression through non-genetic inheritance, which includes parental effects and the transfer of epigenetic factors such as patterns of DNA methylation and histone modification, and microRNA activity (Dai et al. 2020; Spadafora 2020; Venney et al. 2020). TGP can potentially be beneficial when a parent’s environment coincides with what their offspring are expected to experience (Bernatchez 2016; Ashe et al. 2021). For example, TGP could be beneficial in a population that experiences environmental warming over multiple generations whereby the offspring are phenotypically “primed” for a warmer environment (Donelan et al. 2020; McCaw et al. 2020; Venney et al. 2022). In the context of climate change, this phenotypic priming could buffer the effects of rapid warming, allowing additional time for evolutionary processes to occur (Bernatchez 2016; Ashe et al. 2021), though the extent to which TGP can be considered adaptive is arguable (Uller et al. 2013; Sánchez-Tójar et al. 2020). Nevertheless, TGP with temperature has been identified in a number of species (Greenspoon and Spencer 2018; Yin et al. 2019). For example, the coral reef fish *Acanthochromis polyacanthus* exhibits TGP for increased temperatures by improving its aerobic scope in a warm environment by increasing maximum metabolic rate or decreasing resting metabolic rate (Donelson et al. 2012; Bernal et al. 2018). Many of the studies on thermal TGP in aquatic organisms have used eurythermal, temperate or tropical model species; by contrast, cold-adapted aquatic vertebrates are relatively understudied but could stand to benefit from TGP given their vulnerability to environmental warming due to climate change.

Acute and chronic temperature acclimation has obvious impacts on the metabolism of ectotherms, at both the whole organism and molecular level (Pérez-Ruzafa et al. 2018; Petitjean et al. 2019; Morash et al. 2021). Examining the differential gene expression associated with these impacts on metabolism and thermal experiences in general can be informative (Oomen and Hutchings 2017), and some studies have reported the differential gene expression underlying previously observed transgenerational effects (Shama et al. 2016; Bernal et al. 2018; Veilleux et al. 2018). In marine sticklebacks, for example, genes involved in metabolism, mitochondrial protein synthesis, hemostasis and apoptosis were differentially expressed in offspring depending on the multigenerational thermal experiences down the maternal line (Shama et al. 2016). These findings underscore the relationship between the molecular mechanisms underlying physiological plasticity and parental thermal experiences.

We sought to explore differential gene expression in lake trout (*Salvelinus namaycush*) to examine the molecular processes potentially underlying transgenerational responses to elevated temperatures. The lake trout is a long-lived, cold-adapted salmonid that is largely limited to cold, oligotrophic lakes in formerly glaciated regions of North America (Riley et al. 2021; Wilson and Mandrak 2021) and is under significant threat from climate change (Casselman 2008; Guzzo and Blanchfield 2017). Being stenothermal, lake trout behaviourally thermoregulate by migrating to deeper, cooler water when lakes thermally stratify during the summer (Martin and Olver 1980; Guzzo and Blanchfield 2017). Low genetic variation for some lake trout populations (Perrier et al. 2017), and little within- and among-population variation for thermal acclimation capacity (McDermid et al. 2013; Kelly et al. 2014) may limit the adaptive potential of lake trout populations for warming temperate and arctic habitats. A previous study investigated the capacity for transgenerational thermal plasticity in lake trout as a means of coping with warming associated with climate change and found that they exhibit only limited transgenerational plasticity at the whole organism (phenotypic) level (Penney et al. 2021). Unexpectedly, cold-acclimated offspring of warm-acclimated parents exhibited a higher resting metabolic rate than those with one or both cold-acclimated parents (Penney et al. 2021). This study confirmed that the parental environment can influence offspring phenotypic expression, but did not assess potential mechanisms underlying this transgenerational response.

In the current study, we assessed gene expression in previously studied offspring from factorial mating crosses between cold- and warm-acclimated lake trout adults (Penney et al. 2021) to assess transcriptional responses to within- and transgenerational thermal acclimation. Acclimation of the parents (transgenerational) to one of two environmental temperatures (10°C or 17°C) was combined with subsequent acclimation of the offspring (within-generation) to cool (11°C) or warm (15°C) temperatures (Fig. 1). Evidence exists to support both maternal and paternal contributions to transgenerational effects (Marshall 2015; Shama et al. 2016; Penney et al. 2021). Our experimental design allowed us to examine the transgenerational effect of temperature acclimation of each parent, and both parents together, in combination with offspring (within-generation) temperature acclimation on gene expression in juvenile lake trout.

**Figure 1:**
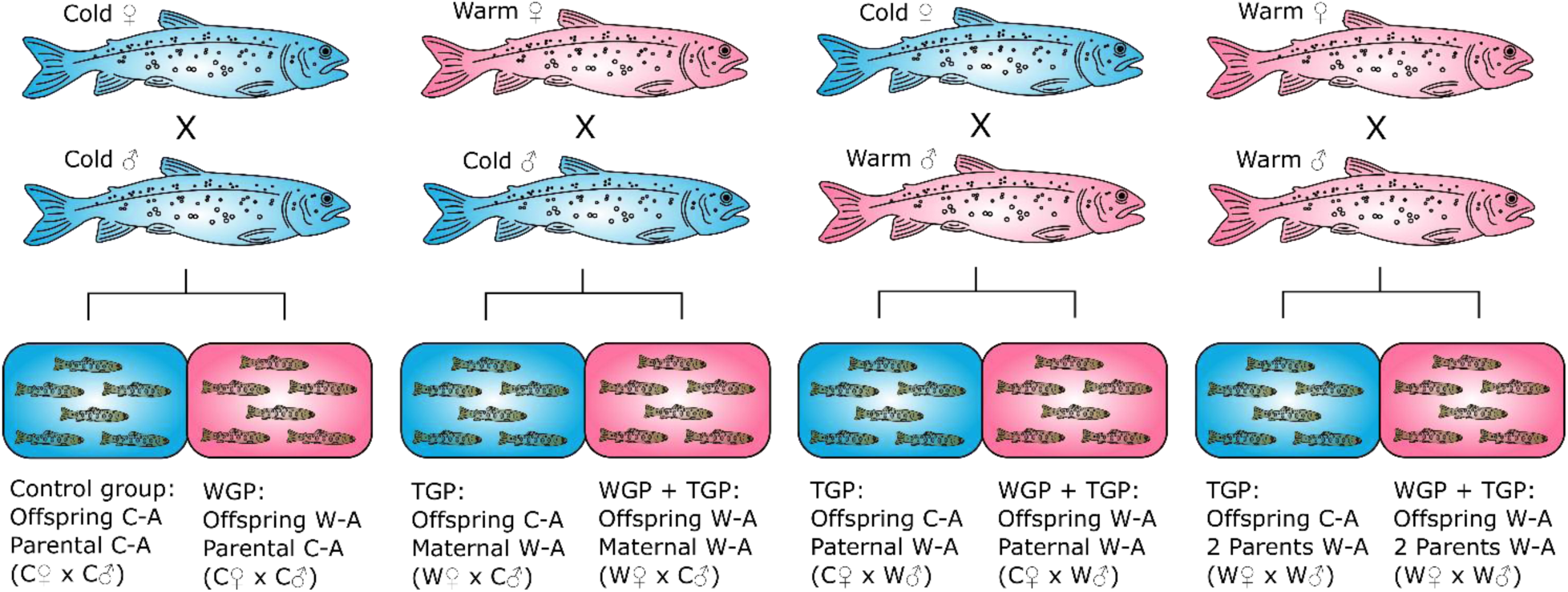
Experimental treatment groups of lake trout (*Salvelinus namaycush*). Two males and two females each from cold- and warm acclimation (C-A and W-A, respectively) tanks were bred in a full factorial cross, resulting in four families per parental treatment group (C_♀_xC_♂_, W_♀_xC_♂_, C_♀_xW_♂_, W_♀_xW_♂_; one each depicted here for simplicity). Offspring from each family was divided and acclimated to either a cold (control; blue: 11°C) or warm (pink: 15°C) temperature.

We hypothesized that genes associated with metabolism, growth and thermal stress/tolerance would be differentially expressed in juvenile lake trout depending on both the acclimation temperature of the offspring and the acclimation temperature of their parents. Based on studies of within-generation and transgenerational warming on differential gene expression in fish (Quinn et al. 2011; Veilleux et al. 2015; Akbarzadeh and Leder 2016; Shama et al. 2016), we predicted that 1) genes involved in growth, heat shock and hypoxia responses, and metabolic pathways would be upregulated in warm-acclimated offspring, and 2) differential expression of these genes would vary depending on whether one or both parents were warm-acclimated, although the size and direction of the effect is difficult to predict given that parental influence on offspring gene expression has been shown to be sex-specific; depending on the sex of the offspring and that of the parent (Best et al. 2018; Baustista et al. 2020).

## Methods

Hatchery-raised adult lake trout from the Ontario provincial Seneca Lake broodstock were used as parents for the experiment. This hatchery strain originated from Seneca Lake, one of the Finger Lakes in central New York State (42°41’ N, 76°54’ W). The Ontario Ministry of Natural Resources (OMNR) hatchery system has maintained this lake trout strain for over five generations, using rotational line crossing (Kincaid 1977) to minimize inbreeding and maintain the founding genetic diversity of the strain (OMNR Fish Culture Stocks Catalogue 2005).

The juvenile (young of year) lake trout used for this experiment were also previously used in a separate, earlier study (Penney et al. 2021). The details of this earlier experiment that are relevant to the current study are summarized in the next section below.

All experiments adhered to the Canadian Council on Animal Care guidelines and were approved by the Trent University Animal Care Committee (Protocol # 24794) and the OMNRF Aquatic Animal Care Committee (Protocol # FACC_136).

### Experimental design and rearing conditions

Mature lake trout (age 8; 2.3 - 4.2 kg) were split into two groups with both sexes kept together (n=8: 4 males and 4 females, and n=9: 4 males and 5 females), and each group was held in a 6,000 L flow-through tank (1 × 1 × 6 m) at the OMNR White Lake Fish Culture Station (Sharbot Lake, Ontario, Canada) with water being supplied from a nearby lake. As the source waterbody (White Lake) began to stratify in early June, 2015, tank water temperatures were increased by 1°C per day starting from approximately 9°C until one tank reached 10 ± 0.5°C and the other 17 ± 0.5°C for the cold-acclimated and warm-acclimated adult lake trout groups, respectively. These temperatures (10°C and 17°C) were chosen because 10°C represents the thermal requirements for lake trout spawning while 17°C was intended to induce physiological thermal stress without preventing reproduction at the higher temperature (Casselman 2008; Penney et al. 2021). Temperatures were controlled and maintained by drawing in and mixing water from above and below the lake’s thermocline as it flowed into the holding tanks. Fish were acclimated to these temperatures for approximately 3 months, mirroring thermal stratification in the source waterbody. Starting in mid-September, the temperature in the warm-water treatment was allowed to gradually cool by holding the proportional inflows from above and below the thermocline constant as the lake’s surface water cooled. Temperatures in the two tanks converged at 10°C after fall turnover (late September) in the lake, and gradually cooled to 7.8°C by the end of the breeding interval (mid-November).

Adult lake trout were reproductive by the end of October with the warm-acclimated fish first spawning on October 30^th^. The cold-acclimated adults were first ready to spawn on November 5^th^ and all mating crosses were completed by November 19th. Experimental offspring families were produced by a full factorial mating cross using two males and two females from each of the two temperature treatments (4 × 4, 8 adults in total) resulting in a total of 16 offspring families; four families from each of four parental treatment groups (W_♀_xW_♂_, W_♀_xC_♂_, C_♀_xW_♂_ and C_♀_xC_♂_, where W = warm-acclimated, and C = cold-acclimated; Fig. 1). Fertilized eggs were transported to the OMNR Codrington Fish Research Facility (Codrington, Ontario, Canada) where they were transferred into 200 L tanks receiving freshwater at ambient temperature (5-6°C) and natural photoperiod under dim light with each family separately contained in steel mesh boxes (9 × 9 × 7.5 cm, one family per box).

In March 2016, when hatched offspring were ready to begin exogenous feeding, we randomly selected 14 individuals from each of the 16 families to transfer to 200 L tanks for temperature acclimation. Seven fry from each family were acclimated to a cold temperature (11°C) and the other 7 from each family were acclimated to a warm temperature (15°C). The offspring’s lower acclimation temperature differs from that of their parents because of reported differences in optimum temperature for both life stages (11 vs 10°C; Edsall 2000, Casselman 2008). The upper acclimation temperature for the offspring represents the anticipated increase in temperature associated with climate change by year 2100 (Hayhoe et al. 2010; IPCC 2021), however, for the parents the upper acclimation temperature was two degrees higher to ensure that temperature was high enough to elicit a physiological thermal response without compromising reproduction. To begin acclimating the offspring, water temperature was increased at a rate of 1°C per day until target temperatures were reached (11°C and 15°C). Due to space constraints, tanks were subdivided by adding steel inserts with perforated bottoms. Each insert was divided into 4 sections (24 × 25 × 28 cm per section) with two paternal half-sibling families (n = 7 each; shared male parent) housed per section. Offspring from pooled families were later individually identified to family using microsatellite genotyping (Penney et al. 2021).

After the 3-4 week acclimation period, offspring were subjected to an acute temperature increase of +2°C·h^-1^ from their acclimation temperature until loss of equilibrium was observed. Eight fish from the same acclimation temperature (11°C or 15°C) were simultaneously tested per day, with each fish held in its own experimental chamber (Penney et al. 2021). Following loss of equilibrium, each fish was quickly transferred to a bath of 0.3 g l−1 of buffered tricaine methanesulfonate (MS-222; Aqua Life, Syndel Laboratories Ltd, BC, Canada) for euthanasia. Whole liver was rapidly dissected from each fish, blotted on a lab wipe and preserved in RNAlater (Invitrogen, Thermo Fisher Scientific) as per manufacturer’s instructions. After 24-48 hours in a 4°C refrigerator, the RNAlater was pipetted from the tissues and the samples were stored at −80°C until RNA extraction. Liver tissue was collected between June 28 and August 9, 2016. We chose to study gene expression in the liver because it is a metabolically active tissue involved in a number of physiological processes that respond to warming (Quinn et al. 2011; Akbarzadeh and Leder 2016; Dammark et al. 2018), and transcriptional responses to multigenerational warming have previously been observed in the livers of zebrafish (Luu et al. 2021).

### RNA isolation and sequencing

RNA was extracted from the preserved liver tissues using a phenol-chloroform extraction method (Chomczynski and Sacchi 2006) after the tissues had been individually homogenized via a FastPrep-24 BeadBeater (MP Biomedicals) with 2 mL Lysing Matrix D tubes (MP Biomedicals) and 1 mL of Trizol reagent (Invitrogen, Thermo Fisher Scientific). RNA was precipitated with RNA precipitation solution (Sambrook and Russel 2001) and isopropanol, and washed with 75% ethanol. RNA samples were resuspended in nuclease-free water (Thermo Fisher Scientific). The purity and concentration of the RNA was quantified using a NanoDrop-8000 spectrophotometer, and RNA quality was assessed by gel electrophoresis of glyoxylated RNA on a 1.5% agarose gel (Sambrook and Russel 2001).

Two lots of RNA samples were sent for high-throughput sequencing over a two-year span. In 2018, liver RNA samples from 24 randomly chosen individuals were sent to The Centre for Applied Genomics (Sick Kids Hospital, Toronto, Ontario, Canada), and in 2020 another 30 liver samples (individuals) were sent to Genome Quebec (Montreal, Quebec, Canada). Altogether, the 54 samples included tissue from six individuals from each of the seven experimental treatment groups, however, the control group (11°C-acclimated offspring from cold-acclimated (CxC) parents) had a total of 12 individuals sequenced: 6 individuals in 2018 and an additional 6 in 2020, so that each treatment group was compared against the control group sequenced in the same year to eliminate batch effects in the analysis of differential expression (Table 1). Individuals from each family were represented in their respective treatment group (there were 4 families per offspring-parent treatment group) and because each sequenced group comprised of 6 individuals, each group had 2 representatives from 2 of the 4 families. Overall, this did not lead to a bias toward any one family being overrepresented in the samples, except for the 2020 control group which had 3 individuals from one family, but this family is not overrepresented in any of the other groups. Subsequent genetic testing for males and females using the salmonid *sdY* sex maker (Yano et al. 2013) confirmed that each group also contained members of both sexes (Table 1). Both facilities assessed the RNA quality via a Bioanalyzer (Agilent Technologies) and all samples passed quality control (RNA integrity number: ≥ 7.5). cDNA libraries were constructed by enriching the poly(A) tails of mRNA with oligo dT-beads using NEBNext Ultra II Directional polyA mRNA Library Prep. In 2018, barcoded libraries were distributed among two and a half lanes and were sequenced on the Illumina HiSeq2500 instrument producing an average of 28 million reads per sample (n = 24, paired-end, 2×100 bp). In 2020, libraries were sequenced on the Illumina HiSeq4000 with barcoded libraries distributed among three lanes producing 36 million reads per sample (n = 30, paired-end, 2×126 bp).

**Table 1:**
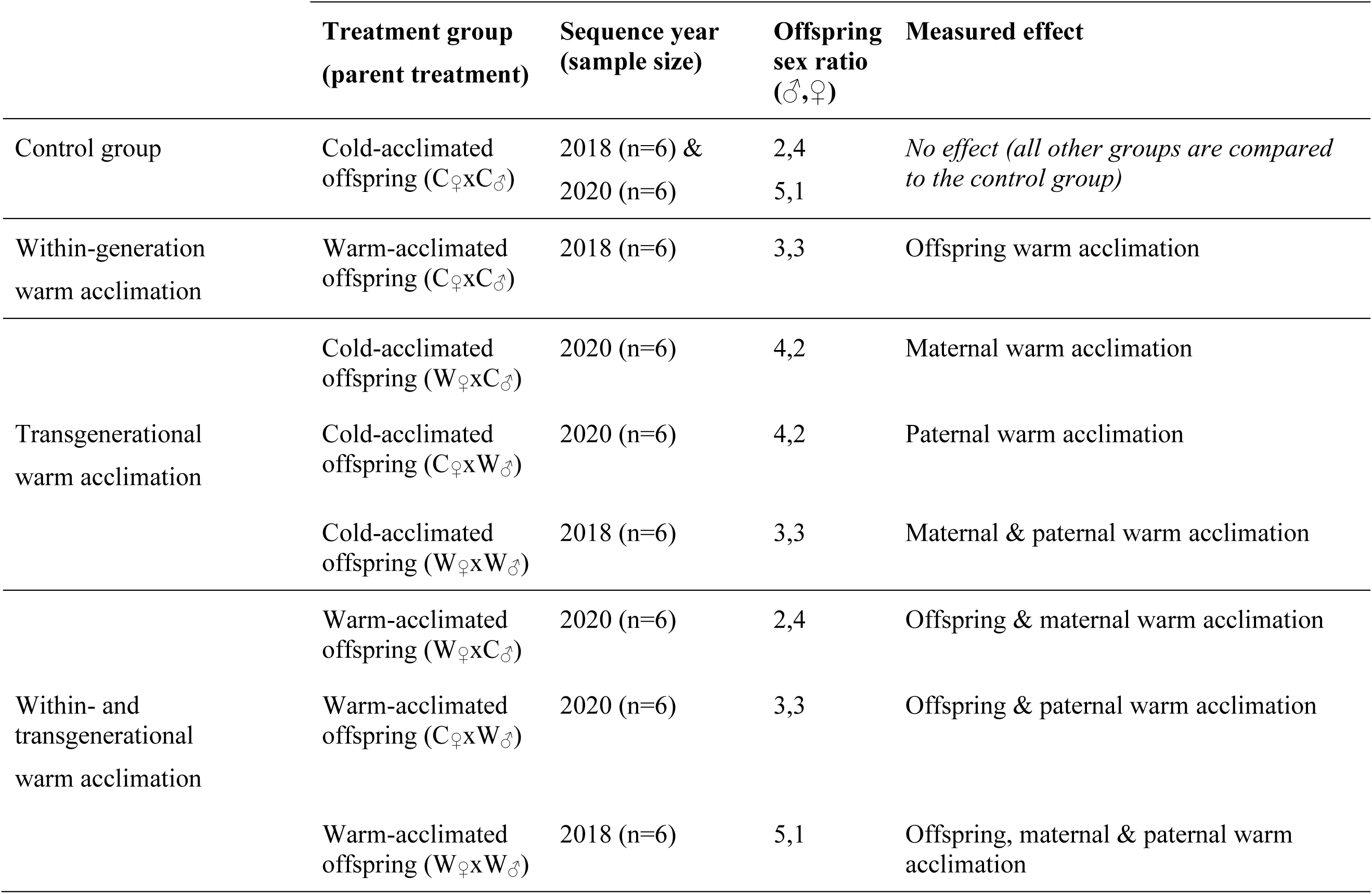
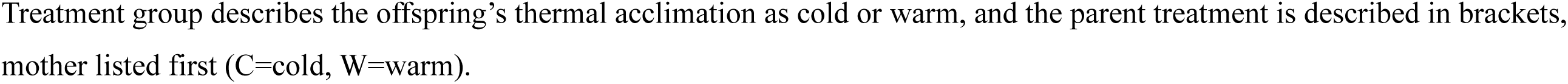
Details for the lake trout offspring groups used to assess differential gene expression following within- and transgenerational thermal acclimation as shown in Figure 1. The measured effect is determined by comparing each treatment group against the control group.

### Data processing and analysis

To analyze reads we used GenPipes, the main in-house framework of the Canadian Centre for Computational Genomics (C3G), to perform major processing steps (Bourgey et al. 2019). Adapters and reads with a Phred quality score of <30 were trimmed using Trimmomatic (v.0.36; Bolger, Lohse and Usadel 2014). The trimmed reads were aligned to the SaNama_1.0, GCF_016432855.1 lake trout (*Salvelinus namaycush*) reference genome (Smith et al. 2022) using STAR (Dobin et al. 2012) and read counts were obtained using HTSeq (Anders et al. 2015).

To measure the effect that offspring (within-generation) and parent (transgenerational, one or both parents) warm acclimation had on gene expression in juvenile lake trout, we compared the differential gene expression of each treatment group to that of the control group. We had a total of seven treatment groups and one control group, providing us with a total of seven comparisons which allowed us to test the effect of offspring (within-generation) warm acclimation in isolation, parent (transgenerational) warm acclimation (either parent in isolation, and both parents together), and the combined effect of the warm acclimation of the offspring with one or both parents. A summary of the comparisons and the effect measured is provided in Table 1. Differential gene expression was determined for each experimental treatment group by comparing it to the control group within a common year to attempt to reduce or eliminate a potential batch effect due to sequencing year in the expression data. For example, experimental groups that were sequenced in 2020 were compared against the control group also sequenced in 2020.

Differences in gene expression levels between the treatment groups were identified using DESeq2 (Love et al. 2014) using negative binomial GLM fitting and Wald statistics (nbiomWaldTest), and ashr (Stephens et al. 2017) was used to shrink log2 fold changes (LFC) in the gene expression data. Significance testing was accomplished by obtaining adjusted *p*-values via the Benjamini-Hochberg approach for detecting false discovery rate (FDR) and differentially expressed genes were considered as those with an adjusted p-value (FDR) of 0.05. Significantly expressed genes with a Log2 Fold Change −0.5 and 0.5 were used in a functional enrichment analysis with Gene Ontology (GO) in clusterProfiler (Yu et al. 2012), using the entire list of expressed genes as background and using the Benjamini-Hochberg method to test for FDR of the enriched terms. We plotted the top five most significantly enriched (i.e. lowest adjusted p-value) GO terms for each category (molecular function: MF; cellular component: CC; and biological process: BP) for each of the seven offspring comparison groups.

A principal component analysis (PCA) was performed using the rlog counts of all genes to explore the variance across all samples and to double check for possible batch effects between the 2018 and 2020 datasets. Seven samples appeared as outliers on the PCA plot. Of these seven samples, two were from the control group (one from 2018 and one from 2020), and a sample each from the following experimental groups: warm-acclimated offspring (C_♀_xC_♂_), cold-acclimated offspring (C_♀_xW_♂_), cold-acclimated (W_♀_xW_♂_), warm-acclimated offspring (W_♀_xC_♂_), and warm-acclimated offspring (W_♀_xW_♂_). To account for the batch effect, we simply matched the samples to the corresponding sequencing group as well as the experimental groups into the differential expression design formula allowing us to account for both batch effect and treatment group to identify differentially expressed genes. We applied the batch correction to the samples using the twelve control group individuals (i.e. n=6 from 2018 and n=6 from 2020 control groups combined) as a shared baseline. One other sample, from a warm-acclimated offspring (W_♀_xW_♂_) experimental treatment group, had a substantial number of reads align to *Homo sapiens* genome indicating potential contamination had occurred during RNA isolation. This sample was excluded from the final dataset.

## Results

The total number of paired-end reads for each of the 53 lake trout liver samples ranged from 26,150,786 to 52,743,330. Between 88.0% and 93.2% of these reads successfully aligned with the lake trout reference genome (Smith et al. 2022).

The PCA of offspring gene expression data did not show clear segregation among sample groups or a significant batch effect between the two sets of sequence data (Fig. 2). Seven samples appeared as outliers on the PCA plot: of these seven samples, two were from the control group (one from 2018 and one from 2020), and single outliers were detected from the warm-acclimated offspring (C_♀_xC_♂_), cold-acclimated offspring (C_♀_xW_♂_), cold-acclimated (W_♀_xW_♂_), warm-acclimated offspring (W_♀_xC_♂_), and warm-acclimated offspring (W_♀_xW_♂_) experimental groups. Before the batch correction was applied, principal components (PC) 1 and 2 explained 26% and 9% of the variance across all groups (Fig. 2A). The variance explained by PC1 and PC2 changed to 11% and 11% respectively after the batch correction was applied (Fig. 2B), suggesting that a batch effect was originally present in our dataset. Other principal components explained 9% or less of the variation (Fig. 2C). While the correction method we used was effective at mitigating the batch effect in our dataset, we are aware that some of the batch effect may still be present to a lesser extent and may influence the interpretation of the results.

**Figure 2:**
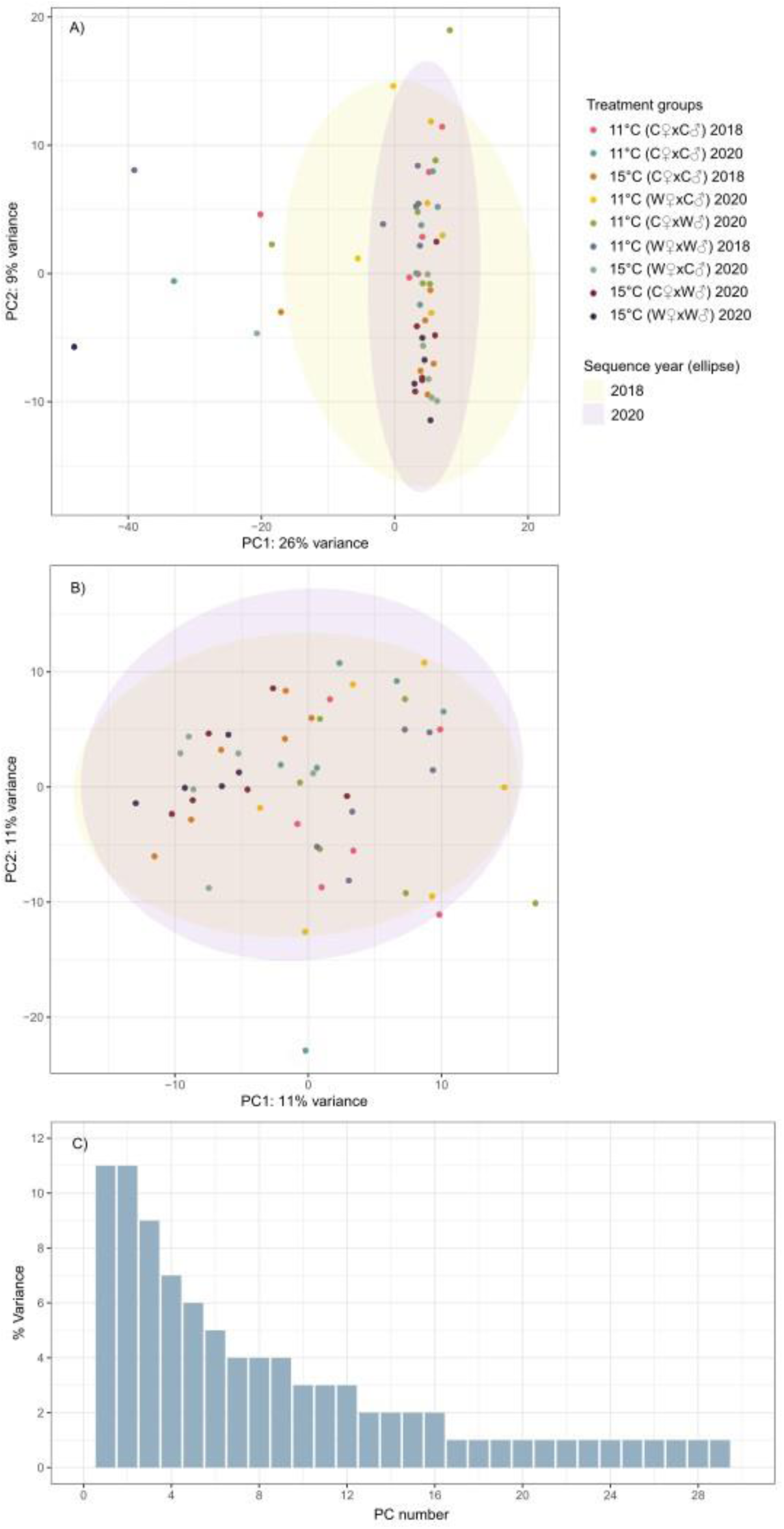
Principal component analysis to explore variance and correct the batch effect between sequencing runs in the gene expression data of the lake trout offspring groups. The PCA plots show ordination of all samples A) before and B) after batch correction. Panel C shows a scree plot of all principal components that cumulatively explain approximately 90% of the variance in the dataset.

### Within-generation acclimation: offspring warm acclimation

Offspring within-generation warm acclimation resulted in 821 DEGs with a LFC of ≤ −0.5 to ≥ 0.5 (Fig. 2 and 3). Of these genes, 318 were uniquely differentially expressed under the isolated effect of within-generation warm acclimation and an additional 106 DEGs were differentially expressed whenever offspring were warm-acclimated (Fig. 3).

**Figure 3:**
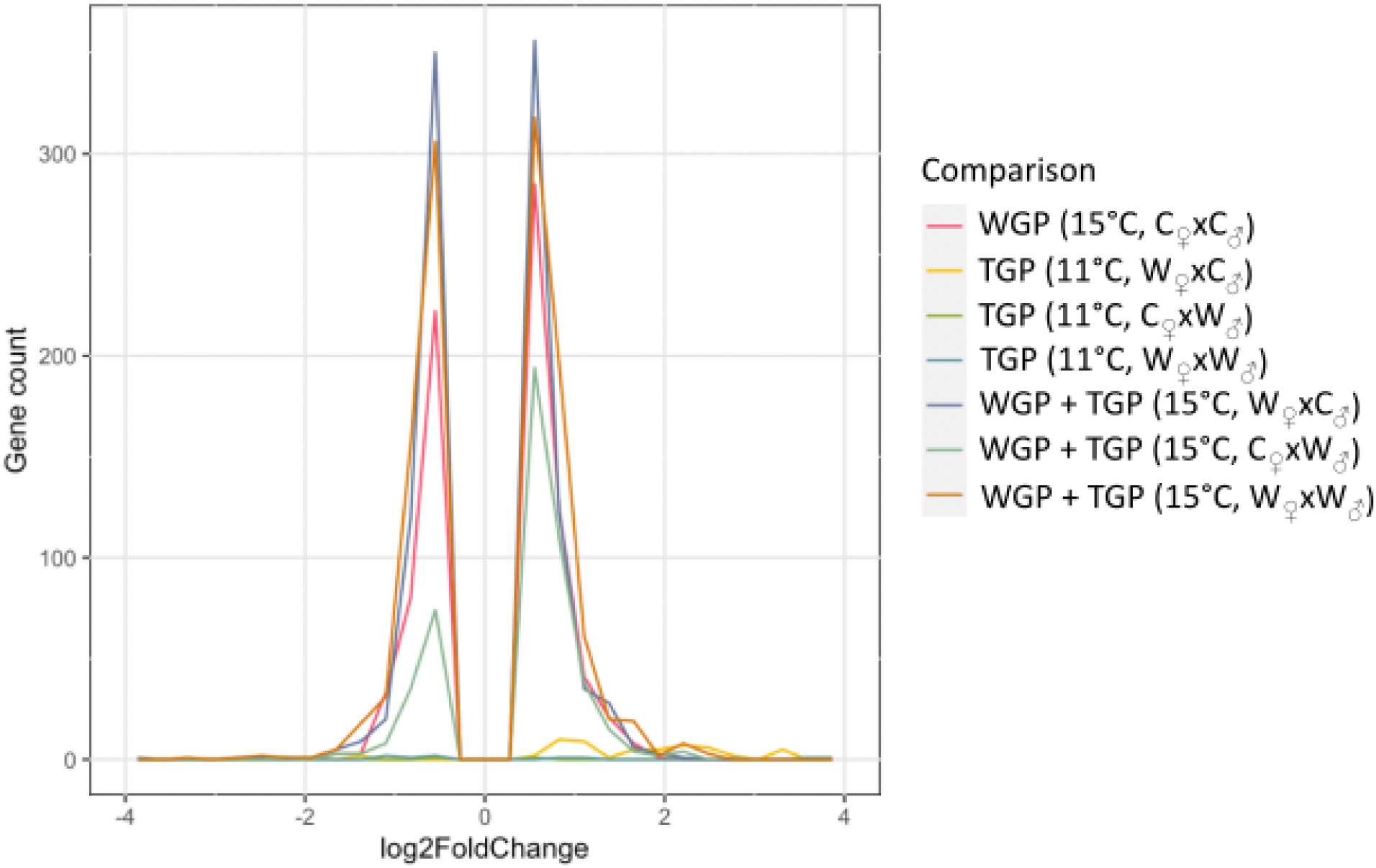
Distribution of up- and downregulated differentially expressed genes in juvenile lake trout with within-generation warm acclimation (WGP), transgenerational warm acclimation (TGP) and the combined effect of both conditions (WGP + TGP) compared against the control group (11°C C_♀_xC_♂_; not shown). Each comparison represents a treatment group (n = 6 individuals). Offspring acclimation temperature (11 or 15°C) and parental treatment (C=cold, W=warm) are provided in parentheses. Gene expression levels are represented as log2-fold change with negative values representing downregulated genes, and vice versa.

The isolated effect of offspring within-generation warm acclimation impacted functions in each GO category (molecular function: MF; cellular component: CC; and biological process: BP) related to oxygen binding and gas transport (GO:0005344, 0020037, 0005833, 0031838, 0072562, 0015669; Fig. 4). Also affected were biological processes and molecular functions related to immune function and cell signaling (GO:0048018, 0030546), such as cellular responses to biotic stimulus/lipopolysaccharide/molecules of bacterial origin (GO:0071222, 0071219, 0071216), and biological processes related to dendrite membranes (GO:0032590) and toll-like receptor signalling (GO:0002224; Fig. 4). Offspring warm acclimation also affected aggresome activity (GO:0016235) and cytokine activity (GO:0005125; Fig. 4).

**Figure 4:**
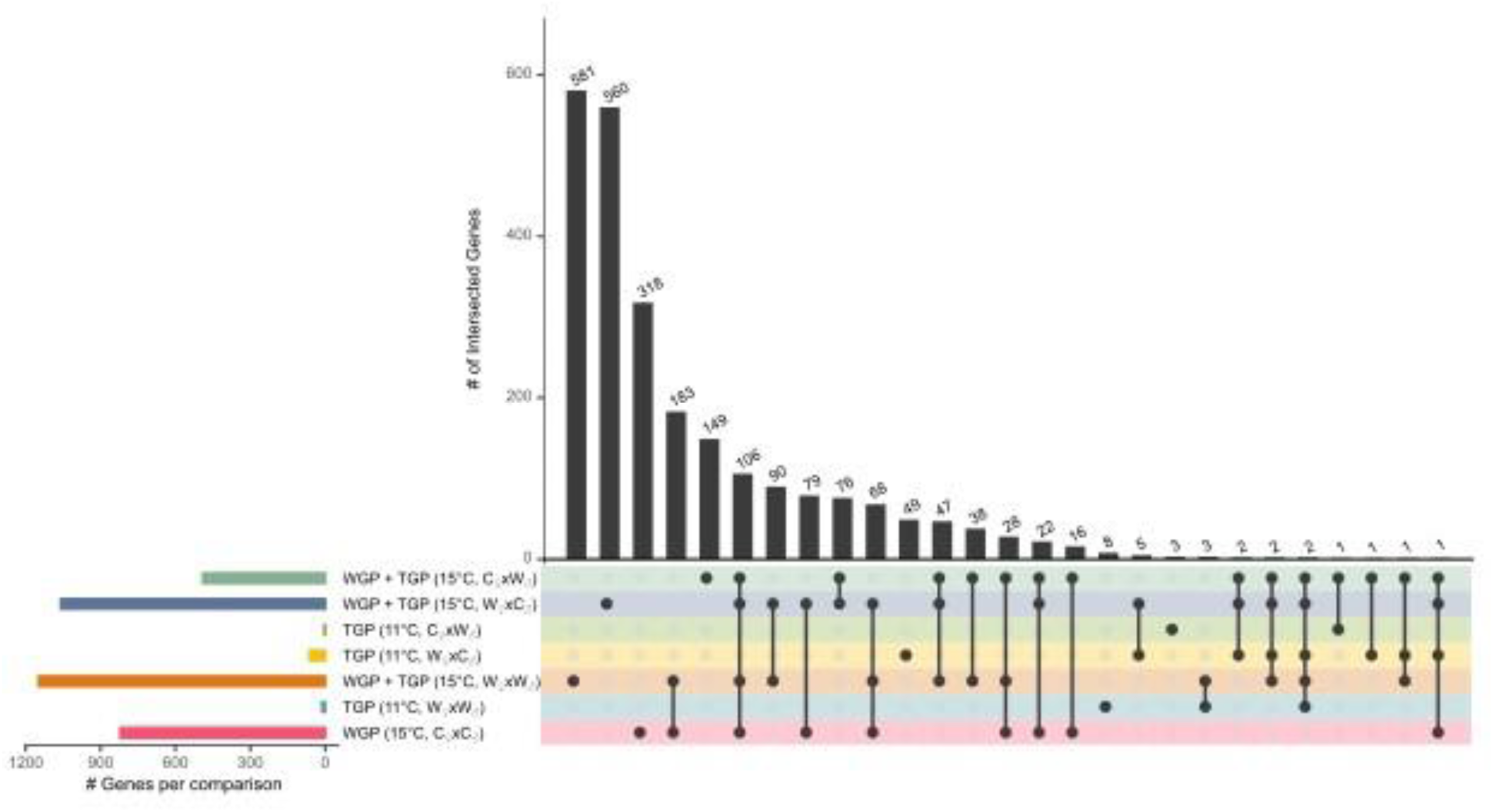
The number of differentially expressed genes that were commonly expressed (or overlapped) between the juvenile lake trout offspring treatment groups (n=6 per group) compared against the control group (11°C C_♀_xC_♂_). The number of differentially expressed genes is listed above each vertical bar. The dots under each bar denote the groups that share those genes (i.e. overlapped genes between groups). A single dot beneath a bar means that these genes are uniquely expressed in that one group. Treatments included within-generation warm acclimation (WGP), transgenerational warm acclimation (TGP) and the combined effect of both conditions (WGP + TGP). Offspring acclimation temperature (11°C or 15°C) and parental treatment (C=cold, W=warm) are provided in parentheses.

### Transgenerational acclimation: parental warm acclimation

An effect of transgenerational (parental) warm acclimation on offspring differential gene expression was evident, however, parental warm acclimation resulted in fewer DEGs compared to the isolated effect of offspring (within-generation) warm acclimation (Fig. 2 and 3). Paternal warm acclimation had the smallest effect, resulting in just 4 DEGs in their offspring (third row down, Fig. 3). The combined effect of warm acclimation of both parents resulted in fewer differentially expressed genes in the offspring compared to when offspring were under the influence of maternal warm acclimation alone (both parents: 13 DEGs vs. maternal: 63 DEGs; sixth and fourth row down, Fig. 3), suggesting that paternal warming can dampen some of the effect of maternal warm acclimation on offspring gene expression. Of these DEGS, 49 were uniquely expressed (i.e. did not overlap with other transgenerational warm acclimation treatment groups) when mothers were warm-acclimated, 3 DEGs were uniquely expressed with paternal warm acclimation and only 8 DEGs were uniquely expressed when both parents were warm-acclimated (Fig. 3).

Maternal warm acclimation impacted the regulation of cellular components and biological processes related to muscle function, specifically cardiac muscle repolarization, in their offspring (GO:0030016, 0043292, 0031674, 0031430, 0008016, 0060307, 0099625, 0099622, 0099623; Fig. 5). Other enriched GO terms included cellular components associated with the apical plasma membrane (GO:0016324), and molecular functions associated with transmembrane transporter and ion channel activity (GO:0015106, 0015276, 0022834, 0015103, 0008509; Fig. 5). With so few DEGs, no GO output was obtained for the transgenerational effect of paternal warm acclimation or warm acclimation of both parents.

**Figure 5:**
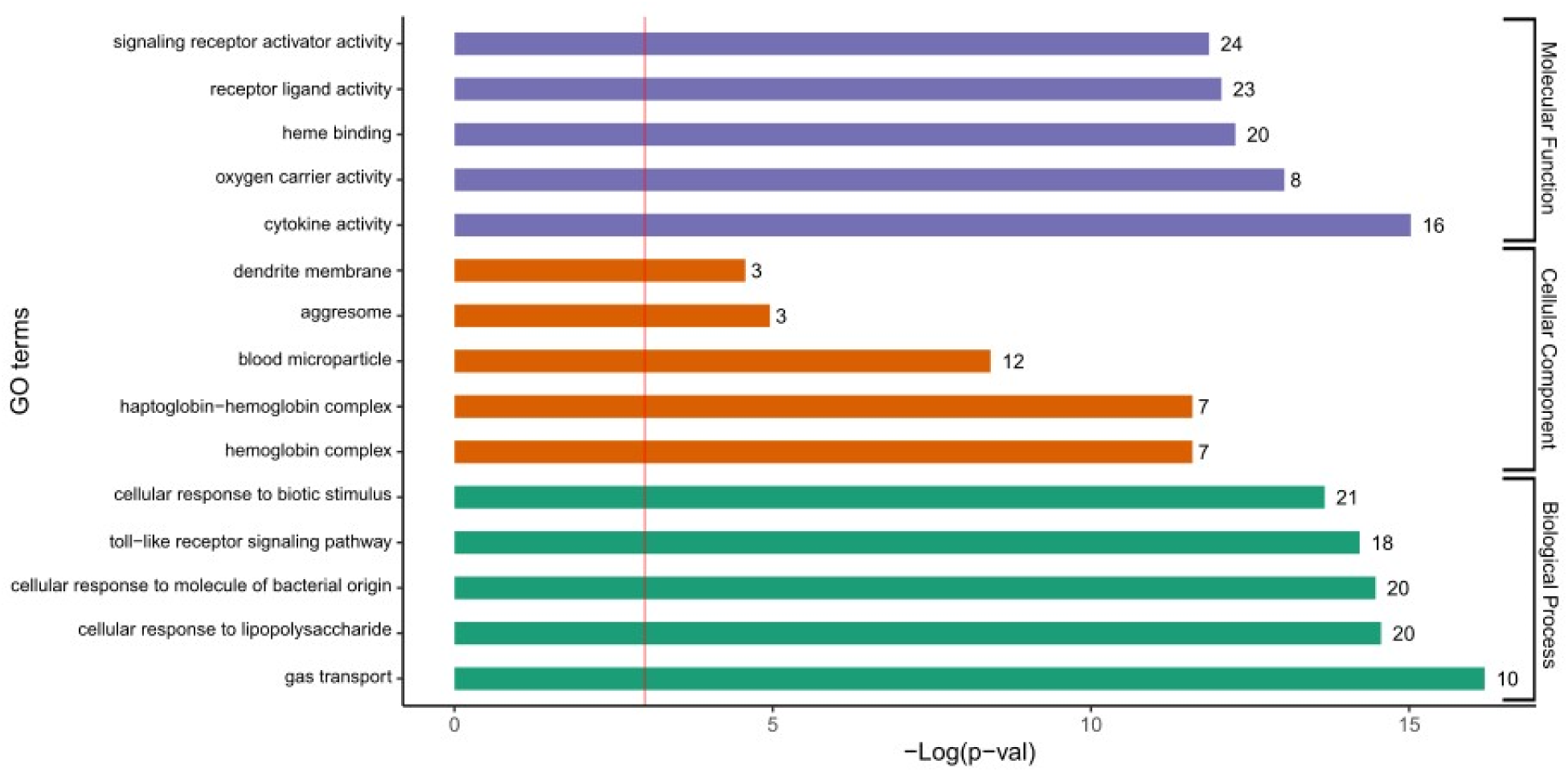
Enriched GO (Gene Ontology) terms for differentially expressed genes in juvenile lake trout offspring (n=6) that were warm-acclimated (within-generation). The GO plot lists the top five most enriched GO terms for each GO category: molecular function (MF), cellular components (CC) and biological processes (BP). The number of differentially expressed genes is provided at the end of each bar. The −log_e_ of the adjusted p-values is given on the x-axis and the threshold (p≤0.05) is indicated by the red vertical line at 2.99.

### Combined generational acclimation: offspring and parental warm acclimation

The combined effect of offspring warm acclimation with that of either or both parents often resulted in a higher number of DEGs than seen with offspring within- or transgenerational warm acclimation in isolation, except for the combined effect of offspring and paternal warm acclimation which resulted in fewer DEGs than seen with the isolated effect of offspring warm acclimation (offspring and maternal: 1,060; offspring and paternal: 492; offspring and both parents: 1,149; Fig. 2 and 3). Unsurprisingly, the greatest number of uniquely expressed DEGs occurred with the combined effect of offspring, maternal and paternal warm acclimation with 581 DEGs (Fig. 3). This was closely followed by the combined effect of offspring and maternal warm acclimation with 560 DEGs, and the combined effect of the offspring and paternal warm acclimation resulted in 149 DEGs (Fig. 3).

When both the offspring and the mothers were warm-acclimated, molecular functions associated with an increased metabolism (e.g. acetyl-CoA C-acyltransferase; GO:0003988) and protein folding chaperone activity (GO:0044183) were enriched (Fig. 6A). Also enriched were molecular functions and biological processes related to DNA synthesis/replication (GO:0004748, 0061731, 0006260, 0006270), and cellular components involving the nuclear membrane activity (GO:0034992, 0034993, 0106083, 0106094; Fig. 6A). Biological processes associated with responses to monoamine catecholamine signaling (e.g. dopamine) were also affected by the combined effect of offspring and maternal warm acclimation (GO:0007191, 0071868, 0071870; Fig. 6A).

**Figure 6:**
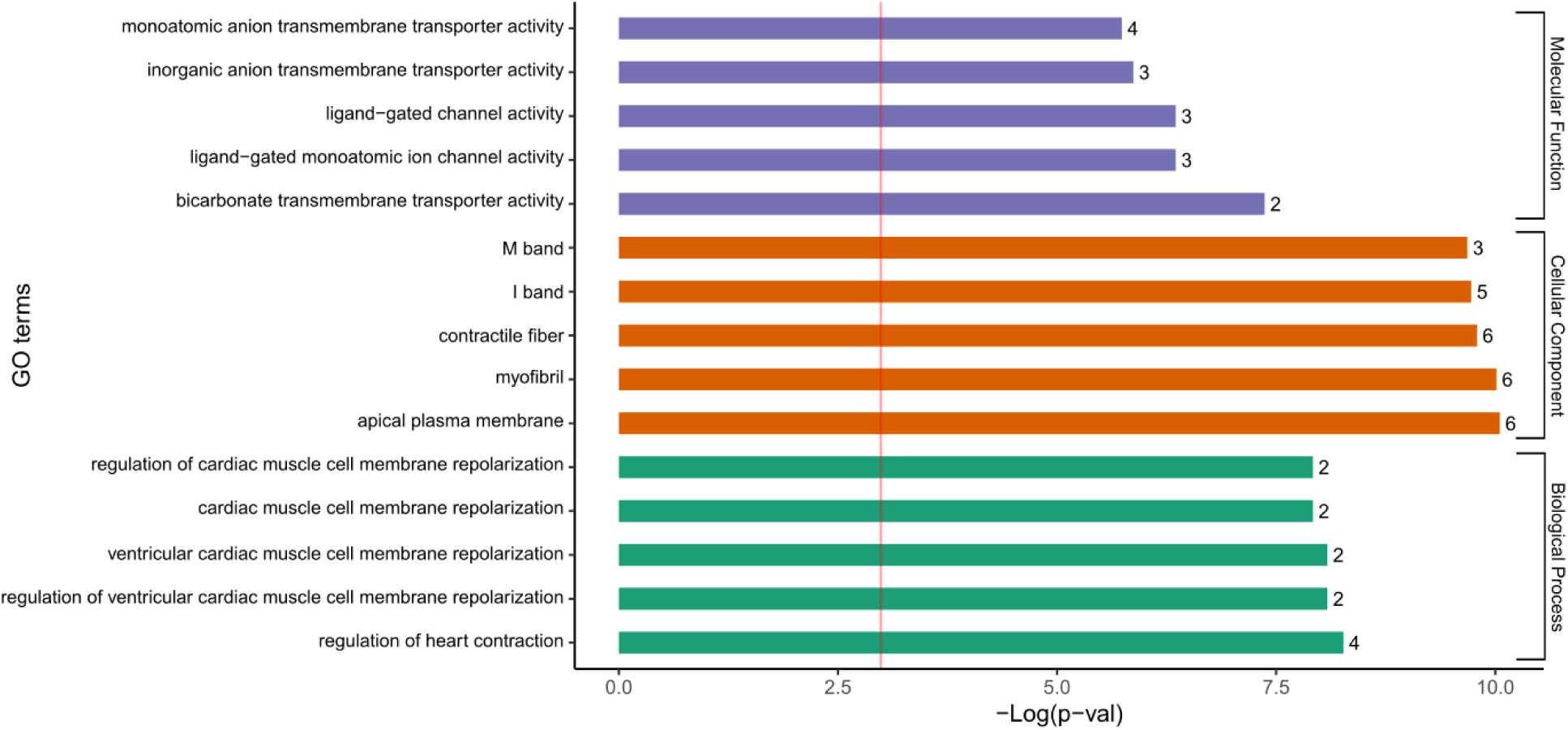
Enriched GO (Gene Ontology) terms for differentially expressed genes in juvenile lake trout offspring (n=6) that received maternal transgenerational warm acclimation. There were no enriched GO terms for paternal transgenerational warm acclimation, or for transgenerational warm acclimation of both parents. The GO plot lists the top five most enriched GO terms for each GO category: molecular function (MF), cellular components (CC) and biological processes (BP). The number of differentially expressed genes is provided at the end of each bar. The −log_e_ of the adjusted p-values is given on the x-axis and the threshold (p≤0.05) is indicated by the red vertical line at 2.99.

**Figure 7:**
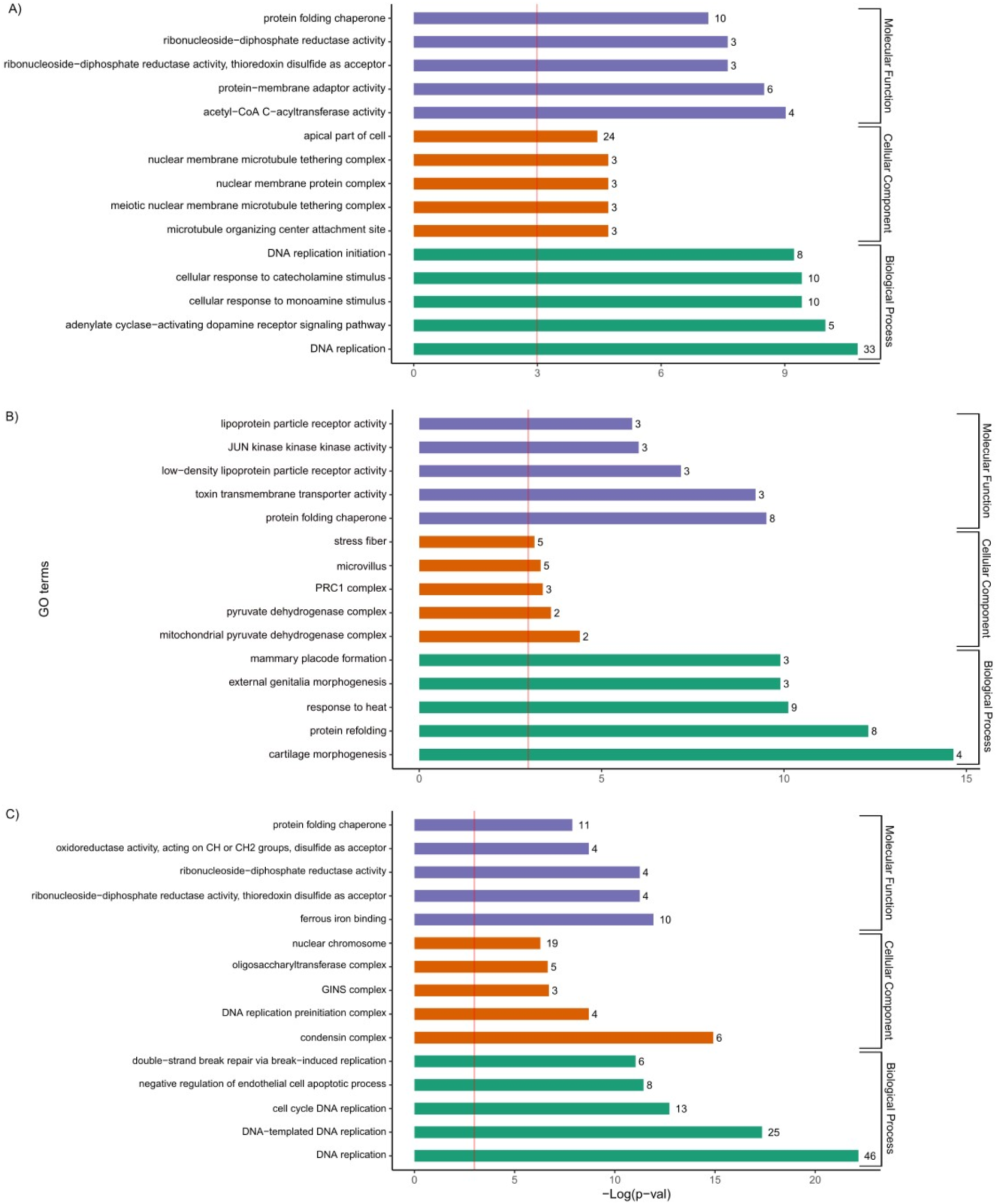
Enriched GO (Gene Ontology) terms for differentially expressed genes in juvenile lake trout offspring (n=6 per group) that were warm-acclimated (within-generation) in combination with transgenerational (parental) warm acclimation of A) the mothers, B) the fathers and C) both parents. The GO plot lists the top five most enriched GO terms for each GO category: molecular function (MF), cellular components (CC) and biological processes (BP). The number of differentially expressed genes is provided at the end of each bar. The −log_e_ of the adjusted p-values is given on the x-axis and the threshold (p≤0.05) is indicated by the red vertical line at 2.99.

When both the offspring and fathers were warm-acclimated (Fig 6B), molecular functions involving protein folding chaperone activity (GO:0044183) and phosphorylation (e.g. JUNKK kinases activity; GO:00004706) were enriched (Fig. 6B). Other molecular functions included membrane receptor and transporter activity for cell uptake of lipoproteins (GO:0019534, 0005041, 0030228; Fig. 6B). Cellular components included increased cellular respiration (e.g. pyruvate dehydrogenase complex activity; GO:0005967, 0045254), DNA transcription (e.g. PRC1; GO;0035102) and components related to the development of microvilli (GO:0005902) and stress fibers (actin filaments; GO:0001725; Fig. 6B). Some enriched biological processes were related to molecular function, such as protein refolding (GO:0042026). Lake trout offspring showed an increased general response to heat (GO:0009408), and indicators of growth, such as cartilage and genital growth (GO: 0060536, 0035261, 0060596; Fig. 6B).

When the offspring and both parents were warm-acclimated, most enriched GO terms across all categories were related to DNA replication and repair (GO:0004748, 0061731, 0016728; 0006260, 0006261, 0044786, 0000727; Fig. 6C). Other molecular functions were related to ferrous binding (GO:0008198) and protein chaperone activity (GO:0044183), while other cellular components included functions involved in protein complexes (GO:0000796, 0031261, 0000811, 0000228), specifically with regards to post-translational modification of protein (e.g. oligosaccharyltransferase complex; GO:0008250; Fig. 6C). Other biological processes were related to apoptotic functions (GO:2000352; Fig. 6C).

## Discussion

Our hypothesis that genes associated with metabolism, growth and thermal stress/tolerance would be differentially expressed in juvenile lake trout depending on the acclimation temperatures of the offspring as well as their parents was supported by the transcriptome data. Genes involved in growth, heat stress responses and metabolic pathways were differentially expressed with within-generation and transgenerational warm acclimation, and with the combination of warm acclimation of both generations. The amount of differential expression (total DEGs) and the type of enriched functions depended on whether it was the offspring or parent(s) that had been warm-acclimated. Within-generation (offspring) warm acclimation had a larger effect on the overall number of DEGs compared to transgenerational contributions from warm-acclimated parents (mother, father or both). As might be expected, however, the combination of offspring, maternal and paternal warm acclimation had the greatest effect on the total number of DEGs.

We considered up- and downregulated differentially expressed genes together in order to assess which pathways/processes are involved in the within- and transgenerational responses to warm acclimation. Although, the both up- and downregulated genes were analyzed together, it should be recognized that warm acclimation often results in disproportionately more upregulated genes compared to downregulated genes (Healy et al. 2017).

### General effect of warming on differential gene expression

In general, within- and transgenerational warm acclimation resulted in differential expression of genes associated with heat tolerance and a heat stress response. In particular, our enrichment analysis highlighted increased aggresome, apoptosis and protein chaperone activity indicating heat-related injury to tissues. Some of these pathways included genes that encode heat shock proteins (HSPs), which is not surprising given that induction of HSPs are evolutionarily conserved across numerous taxa (Krens et al. 2006; Kostenko et al. 2011) and that increased expression of these proteins have been observed previously in other fish taxa following heat stress (Keller et al. 2008; Kelly et al. 2018; Liu et al. 2021). Warm-acclimated lake trout offspring also showed an increase in immune system activity. The lake trout offspring in our study demonstrated differential expression of genes associated with a toll-like receptor signalling pathway, and similar results have been observed in other fish species. For example, upregulation of the toll-like receptor 3 gene (*tlr3*) occurred in the liver of the flounder, *Paralichthys olivaceus*, following heat shock (Lee et al. 2023). An increase in immune-related functions is common in heat stressed fish and often appear to coincide with indicators of temperature-induced tissue damage (Huang et al. 2018; Beemelmanns et al. 2021). Taken together, these pathways indicate increasing temperatures elicited a stress response in lake trout, which we would expect given that each offspring was subjected to an acute temperature increase (+2°C/h) and these pathways were likely upregulated to respond to cellular/protein damage occurring at their upper critical thermal limit (Penney et al. 2021).

With warm acclimation, there was a general enrichment of pathways associated with an increased metabolic demand, including O_2_ binding and gas transport. Upregulation of processes related to O_2_ transport, such as heme binding, have previously been observed in the liver of fish with a history of heat stress (Duan et al. 2024) and could mean that the lake trout offspring were attempting to increase their O_2_ carrying capacity to meet increased metabolic demands (Lu et al. 2020). We also observed differential expression of metabolic enzymes such as acetyl-CoA C-acyltransferase activity and pyruvate dehydrogenase. Both of these enzymes are used in the production of acetyl CoA: a key coenzyme involved in several metabolic functions, including the citric acid cycle (Shi and Tu 2015). Our transcriptomic findings here are indicative of an increase in O_2_ demand which aligns with the findings of Penney et al. 2021 in that offspring metabolic rates, measured as rates of O_2_ consumption (MO_2_), were elevated at the time of tissue sampling.

We found some indication that warm acclimation and upper thermal stress affected the growth of the lake trout offspring. We expected to see differential expression of some growth-related functions given that the warm-acclimated offspring were heavier than the cold-acclimated offspring at the time of sampling (Penney et al. 2021). These functions included oxygen transport, cartilage growth and synthesis of cellular membrane and cytoskeletal components. Other studies have observed upregulation of similar pathways involved in growth in fish (Danzmann et al. 2016; Lu et al. 2020; Vergauwen et al. 2010). For example, we saw an enrichment of acetyl-CoA C-acyltransferase activity which could indicate a higher demand for acetyl-CoA production for growth, just as a transcriptome analysis of the liver of *Salmo salar* revealed an increased demand for acetyl-CoA in faster growing groups of fish (Caballero-Solares et al. 2018).

It is curious that enrichment of more growth-related processes was not discovered in the heavier, warm-acclimated lake trout offspring in our study. For this reason, we cannot be entirely confident that the enriched pathways we observed resulted primarily from growth of the offspring. We collected liver samples at each fish’s critical thermal maximum, so it is possible that fish were responding to the stress of the acute temperature increase and may have suppressed growth during the acute temperature challenge in favour of a heat stress response. Suppression of growth at high temperatures is not unusual. For example, increased expression of the genes *GH* and *IGF-1* (indicators of growth) occurred in *Labeo rohita* at slightly warmer acclimation temperatures but was suppressed at higher acclimation temperatures (Shahjahan et al. 2021). Alternatively, there is evidence to suggest that growth hormone signaling pathways are more sensitive to elevated temperature in the muscle compared to other tissues (Hevrøy et al. 2015). For example, Houde et al. (2019) did not see differential expression of insulin-like growth factor binding proteins *IGFBP-1* and *IGFBP-10-v1* in the gills of *Onchorhynchus tshawytscha*. Thus, it is possible that we may have seen a stronger transcriptional response for growth-related pathways in the muscle rather than in the liver of the lake trout used in our study, though confirmation of this would require further experimentation.

### Transcriptomic responses to transgenerational warm acclimation

We found that an offspring’s thermal acclimation often had a greater influence on the number of DEGs than did the thermal experience of its parents, suggesting that transgenerational transcriptomic responses to warming are weaker than those that occur within a generation. This result agrees with a previous study that saw that the effect of transgenerational warm acclimation on offspring metabolic rate (rate of O_2_ consumption) was limited (Penney et al. 2021). In contrast to our findings, some previously published evidence suggests that transgenerational warming has a stronger effect on offspring differential gene expression compared to developmental warming (Veilleux et al. 2015; Shama et al. 2016). It has also been suggested that transgenerational effects are strongest in early-juvenile life stages (Yin et al. 2019) and it would be interesting to see if this is reflected at the transcriptomics level by looking at gene expression at different life stages following within- and transgenerational warm acclimation in lake trout. Related to this, it may worth noting that prior to warm acclimation all offspring in our experiment were reared in the same environment and temperature regime to minimize differential developmental effects. How developmental and transgenerational warming interact in these fish is not fully understood, but the influence of thermal acclimation on differential gene expression during early, developmental life stages compared to later life stages may be a fruitful area for study. Lastly, the combination of offspring, maternal and paternal warm acclimation had the greatest effect on the overall number genes that were differentially expressed. This adds to the growing literature suggesting that the strength of transgenerational plasticity is greatest when offspring and parental thermal experiences coincided (Shama et al. 2014; Donelson et al. 2017; Yin et al. 2019).

Paternal transgenerational warm acclimation appeared to dampen the effect of maternal warm acclimation on offspring gene expression in that fewer genes were differentially expressed in the offspring when both parents were warm-acclimated compared to when offspring were under the influence of maternal warm acclimation only. Earlier discussions of transgenerational plasticity speculated that fathers could influence maternal investment in their offspring through non-genetic means, including imprinting and components included in glandular secretions (Uller 2008; Bonduriansky and Day 2009). Investigations of paternal effects are less common than that of maternal effects and there have been calls for more studies on paternally mediated non-genetic inheritance on offspring phenotypes (Rutkowska et al. 2020). Recent studies have shown that a stronger paternal contribution is possible. In brook trout (*S. fontinalis*), paternal transgenerational warming can contribute more to offspring thermal responses and survival compared to the maternal contribution (Penney et al. 2022; Houle et al. 2023). In contrast, we did not previously see a stronger paternal effect in the thermal response of the lake trout offspring used for this experiment (Penney et al. 2021). Whether this was due to suppression of offspring gene expression is not entirely clear but further investigation into this effect could be insightful, potentially revealing a mechanism by which fathers can control their offspring’s developmental rate or resource use in competition or in concert with maternal effects.

An effect of parental (transgenerational) warm acclimation on the offspring’s heat shock response was evident, providing support for our hypothesis. The combination of offspring and maternal warm acclimation led to the differential expression of genes associated with cellular responses to catecholamines (e.g. dopamine). Modification of cellular sensitivity to catecholamines can influence responses to warmer temperatures. For example, sensitivity to dopamine can regulate thermal preference behaviour (Ueno et al. 2012), and blocking of the β-adrenergic pathway can lead to a decreased cellular heat shock response in rainbow trout (Templeman et al. 2014). Additionally, biological processes related to protein refolding and a general response to heat were differentially expressed under conditions of offspring and paternal warm acclimation. Our results suggest that perhaps thermal sensitivity is regulated when offspring and parental warm acclimation coincide which is supported by the idea that transgenerational plasticity can be adaptive when environmental changes trend in the same direction across generations (Shama et al. 2014; Donelson et al. 2017; Yin et al. 2019).

Our prediction that transgenerational warm acclimation would upregulate metabolic processes in lake trout offspring was partially supported, and is indicative of a higher metabolic rate in warm-acclimated offspring with warm-acclimated parents. Maternal transgenerational warm acclimation in isolation led to the enrichment of processes related to cellular transmembrane ion transport, including the transfer of bicarbonate ions. The increase in higher metabolic rates of ectotherms with warming can result in metabolic (lactic) acidosis and the release of bicarbonate ions into the blood counteracts the shift in pH (Stewart et al. 2019). The combined effect of warm-acclimated mothers and offspring led to differential expression of genes associated with acetyl-CoA C-acyltransferase activity, and the combined effect of warm-acclimation of the fathers and offspring led to differential expression of genes associated with the pyruvate dehydrogenase complex. These two enzymes are involved in regulating metabolism through their involvement in the production of acetyl CoA (Shi and Tu 2015), however, previous work showed that neither resting or peak metabolic rates differed between warm-acclimated offspring from warm- or cold-acclimated parents (Penney et al. 2021). It is worth noting that gene expression does not necessarily confer a protein to function given that post-transcriptional and post-translational activity could also tag a protein for degradation; the enrichment analysis may suggest an upregulation of metabolic processes, but these proteins may have been rendered inactive, or perhaps were not expressed highly enough to be detected at the level of the phenotype in the lake trout offspring.

We found some support for a transgenerational effect of warm acclimation on growth in lake trout at the transcriptome level. Molecular functions including DNA synthesis (e.g. ribonucleoside-diphosphate reductase activity), biological processes related to DNA replication, and cellular components involved in stabilizing the nuclear membrane were influenced by the combined effect of maternal and offspring warm acclimation. An upregulation in cellular structural components has previously been observed with warm acclimation and growth in fish (Vergauwen et al. 2010; Lu et al. 2020). Protein folding chaperone activity also appeared in the molecular function GO category for each of the combined warm acclimation groups: warm-acclimated offspring with warm-acclimated mothers, fathers, or both parents and may be upregulated for the formation of new proteins. An alternative explanation could be that these functions were involved in other processes related to cell repair/replacement as cells and proteins sustain heat damage, especially considering that apoptosis appeared in the enrichment analysis when offspring and both parents were warm-acclimated. Finally, biological processes related to genitalia and cartilage differentiation were differentially expressed when fathers and offspring were warm-acclimated and may be indicative of growth of reproductive organs as the offspring develop towards sexual maturity. Transgenerational warming can result in larger offspring (Salinas et al. 2012), however, our transcriptome findings are not consistent with the findings of Penney et al. (2021) where transgenerational warm acclimation had no effect on the mass of the offspring. Nevertheless, our transcriptome results suggest that there is some transgenerational plasticity in growth-related functions when both parents and offspring experience warming.

Strangely, warm-acclimated mothers seemed to have an influence over processes related to muscle function in their offspring. These included cellular components and biological processes associated with contractile fibers and cardiac muscle cell membrane function. These same functions were not apparent with the combined effect of maternal and offspring warm acclimation. It is curious that GO terms specifying muscle functions appeared in our results given that the focal tissue was liver. However, a transgenerational effect of thermal acclimation on muscle function has been observed in liver samples fish. For example, transgenerational warm acclimation resulted in differentially methylated regions of genes associated with cardiac muscle function, heart rate and cardiac myofibril assembly in liver tissue of *Acanthochromis polyacanthus* (Ryu et al. 2020). Future studies could test the tissue-specific transcriptomic responses to transgenerational warm acclimation in lake trout or other salmonids to better understand the functions under the influence of transgenerational plasticity.

Venney et al. (2022) used a similar breeding design as this study to assess temperature-related DNA methylation in adult brook trout and their offspring. In contrast to our findings, Venney et al. (2022) observed a stronger transgenerational effect from adults held at higher temperatures during gonad development compared with methylation from offspring rearing temperatures. Despite the apparent contrasts between our results and those of Venney et al. (2022), the findings from the two studies are in fact complementary rather than contradictory. Some studies on salmonids have shown that changes in global methylation do not appear to influence differential gene expression (Christensen et al. 2021; Leitwein et al. 2022). DNA methylation can influence gene expression in an individual and subsequent generations, but gene expression is also a direct response to physiological, metabolic, and bioenergetic requirements including acclimation. The greater methylation observed from reproductive adults by Venney et al. (2022) may reflect the lower temperature tolerances of reproductive adults compared with other free-swimming life stages (Dahlke et al. 2020). The lesser extent of DNA methylation observed in juvenile brook trout by Venney et al. may also reflect the ability of subadult brook trout to use warmer temperatures compared to adults (Smith and Ridgway 2019), as well as the greater metabolic scope of juvenile fish compared with reproductive adults (Dahlke et al. 2020). By contrast, rather than studying methylation, our study examined levels of gene expression and showed a greater effect of within-generation acclimation. Our current results parallel those from a previous study looking at whole-organism respirometry in lake trout (Penney at al. 2021).

### Conclusions

Our study has shown that lake trout exhibit both transgenerational (both maternal and paternal) and within-generation thermal acclimation, with both levels of acclimation influencing the strength of differential expression of genes associated with thermal stress/tolerance, metabolism and growth. Despite this, the limited transgenerational effect on gene expression indicates that adaptive transgenerational acclimation is unlikely to contribute sufficiently substantive benefits to enable lake trout to cope with rapidly changing environmental conditions related to global warming (Penney et al. 2021). Instead, population-level responses to temperature-related stress will likely be restricted to within-generation acclimation (Kelly et al. 2014) and longer-term adaptive responses. Beyond acclimation, persistence of cold-water species and populations may be reliant on existing genetic resources to cope with chronic warming (Comte and Olden 2017; Crozier and Hutchings 2014; Guzzo and Blanchfield 2017), which is particularly daunting for long-lived species (Willi et al. 2006). It therefore seems likely that lake trout and similarly vulnerable cold-water species may require increased management and conservation efforts to ensure their future in a rapidly warming world.

## Acknowledgements

The Ontario Ministry of Natural Resources (OMNR) Fish Culture Section provided adult lake trout, rearing space and logistical support at the White Lake Fish Culture Station. Bill Sloan, Scott Ferguson, Anne McCarthy (OMNR) and John Dewart provided assistance with breeding adult fish; Bill Sloan and Scott Ferguson also provided invaluable logistical support for all aspects of fish husbandry from care of the juvenile trout to assisting with experiments at the OMNR Codrington Fisheries Research Facility. Caleigh Smith (OMNRF) genotyped adult and juvenile fish for parentage assignment, family identification of juveniles and sex marker typing. Paul Craig and Heather Ikert (University of Waterloo) kindly provided advice and guidance on the enrichment analysis. Paul Craig, Graham Scott, Anne Dalziel and Aaron Shafer provided feedback on early drafts of the manuscript. The RNA sequencing was conducted by The Centre for Applied Genomics (Sick Kids Hospital, Toronto, Ontario, Canada) and Genome Quebec (Montreal, Quebec, Canada). The bioinformatic analyses was performed by Canadian Center for Computational Genomics (Montreal, Quebec, Canada). The Canadian Center for Computational Genomics (C3G) is a Genomics Technology Platform (GTP) supported by the Canadian Government through Genome Canada.

Research funding was provided by the Canada-Ontario Great Lakes Agreement (COA) to CCW, by the Ontario Federation of Anglers and Hunters (OFAH) to CMP, and a Natural Sciences and Engineering Research Council (NSERC) Discovery Grant to GB.

## Data archiving statement

Data for this study are available at: *to be completed after manuscript is accepted for publication*.

## Authors’ contributions

CMP, GB and CCW conceived the ideas and designed methodology; CMP collected and analysed the data, and led the writing of the manuscript. FL and GZ completed the bioinformatics. All authors gave final approval for publication.

